# Quantifying genetic, environmental and individual stochastic variability in *Plantago lanceolata*

**DOI:** 10.1101/270603

**Authors:** Ulrich K. Steiner, Shripad Tuljapurkar, Deborah A. Roach

## Abstract

Predicting ecological and evolutionary population dynamics requires understanding how genetic and environmental parameters influence variation in survival and reproduction among individuals. However such a focus often neglects the stochastic events that individuals experience throughout their lives that also influence survival and reproduction. With an illustrative example, we quantify and illustrate the influence of such non-selective demographic variability on population dynamics using size-structured matrix models of an experimental population of *Plantago lanceolata*. Our analysis shows that variation in survival and reproduction among individuals explained by environment, genes, and their interaction was modest compared to the stochastic variation in lifespan and reproduction. We illustrate how expectations on population growth, based on expected lifetime reproduction and generation time, can be misleading when variance in reproduction among individuals of the same genotype (full sibs) was large. Such large within genotype variance can lower population growth, fitness. Our results accompany recent investigations that call for more focus on stochastic variation in survival and reproduction, rather than dismissal of this variation as uninformative noise.

---

Population dynamics are driven by simple demographic events, the survival and reproduction of individuals in a population (Metcalf and Pavard 2007). To understand such dynamics it is important to quantify the sources of variability in vital rates in order to predict evolutionary and ecological change, and to forecast population responses to environmental change. Individual variability in survival and reproduction has been ascribed to genetic variability, environmental stochasticity, and demographic stochasticity (Roughgarden 1975; May 2001; Lande et al. 2003), but these terms rarely differentiate among selective and non-selective, i.e. neutral, causes of variation (Engen et al. 1998). “Genetic variability” might be additive genetic variance, or include non-additive genetic variation and genotype-environment interactions. “Environmental stochasticity” usually includes temporal fluctuations affecting all individuals in a population, but might not include differences below the population level, e.g., as caused by temporal or spatial environmental differences within the population (Lande et al. 2003; Melbourne and Hastings 2008). Additionally, where genotypes are not known, environmental stochasticity frequently includes genotype-environment interactions (Lande et al. 2003). “Demographic stochasticity” is defined by Lande et al. (2003) and Melbourne and Hastings (2008) as the result of independent chance events, e.g., individuals with identical vital rates may differ in how many offspring they produce or how long they live. Demographic stochasticity has also been used to describe demographic heterogeneity, when individuals have distinct probability distributions for reproduction and survival (Melbourne and Hastings 2008). Such differences can be fixed at birth or can change dynamically during the life course (Tuljapurkar et al. 2009), and may in turn be influenced by genetic differences, maternal differences or even chance (Melbourne and Hastings 2008). The amount of demographic stochasticity can be estimated by calculating demographic variance (Vindenes and Engen 2017), a quantity that has earlier been called individual stochasticity (Caswell 2009), and dynamic heterogeneity (Tuljapurkar et al. 2009; Steiner and Tuljapurkar 2012). In spite of these ambiguities, we know that including or excluding different types of variation can substantially affect ecological and evolutionary dynamics (Melbourne and Hastings 2008; Steiner and Tuljapurkar 2012; Vindenes and Langangen 2015; Vindenes and Engen 2017).

In our view, an accounting for the causes of variation in survival and reproduction should focus on the difference between adaptive and neutral variation. As pointed out above, that differentiation is sometimes lost when using the terms genetic variability, environmental stochasticity and demographic stochasticity. Here we consider only two sources of selective variation: genetic variation (additive and non-additive), and variance in phenotypic plasticity (we ignore neutral genetic variation). Hereafter we call these, respectively, genotypic (G) variation and genotype by environment (GxE) variation. We also consider three sources of non-selective variation. Firstly, environmental variation that affects every individual’s vital rates (often defined as environmental stochasticity). Secondly, non-selective demographic variation, stochastic demographic processes that determine the reproduction and survival of an individual and are selectively neutral (Caswell 2009; Tuljapurkar et al. 2009; Steiner et al. 2010; Steiner and Tuljapurkar 2012). Finally, we consider non-selective variation that is true noise due to, e.g., non-selective measurement error, data processing error, or hidden and uncontrolled experimental conditions.

An ideal experiment to quantify, in nature, the sources of variability in survival and reproduction would track many individuals with known genotypes over their lives, with the environment perfectly known, and no measurement or processing errors. Of course such ideal conditions cannot exist. To approach such conditions an increasing number of studies have followed marked and genotyped individuals in the wild (Pajunen VI and Pajunen I 2003; Roach and Gampe 2004; Ozgul et al. 2009; Pennisi 2012). Most of these studies lack the large numbers of individuals needed to reveal causation beyond interpreting a simple correlation between the environment, the genotype, and an individual fitness component (Endler 1986). The limited population sizes and small numbers within pedigrees or genotypes leads to biased correlations and variance decomposition (Hadfield et al. 2010).

Here, as an illustrative example, we analyze data from an experiment in which large numbers of seedlings of the common ribwort plantain, *Plantago lanceolata,* from multiple experimental crosses, were planted in a multiple cohort design and individuals were followed until death. To this end we have many individuals of the same genotype (known crosses) that experience similar environments among years. To decompose the variability in survival and reproduction among individuals we used a matrix model approach to analyze genotype-environment specific population dynamics. For each year-sire combination, we constructed a stage-structured matrix model with size (number of leaves) as the stage character and the number of reproductive stalks (inflorescences) as a measure of reproduction. With each year-sire model we computed macroscopic demographic parameters (population growth rate, generation time, average lifetime reproduction, life expectancy), and the variance in lifespan and lifetime reproductive success. We quantified genotypic variation (G) in lifespan and reproduction by estimating the variance among the 16 sires used for the crosses, which was corrected for the dam effects. We computed the contribution of the genotype by environment interaction (GxE) by the variance among year-sire combinations, again corrected for the dam effects. We used the variation among the six years to determine the non-selective environmental variation, and the variation among 17 spatial blocks to determine small-scale spatial environmental variation. The non-selective demographic variation was estimated directly from the matrix model for any year-sire class. To quantify this variation directly from the model we used the large variation in growth and shrinkage among individuals of the same year-sire combination that went beyond the expected (mean) growth. We assumed that this additional variance in growth (size) was directly or indirectly related to the differences in survival and reproduction among individuals. The stochastic nature of these stage-structured Markov process based models allowed us to directly estimate the non-selective demographic variation in fitness components (Steiner and Tuljapurkar 2012; Steiner et al. 2014) instead of inferring this variation from the residual variances of linear models as for instance done in most quantitative genetic models. Our aim was to disentangle the genetic, environmental and demographic causes of variability among individuals in reproduction and survival and thereby identify drivers of population dynamics.

We find that, despite substantial fluctuation in vital rates among years, the variation in survival and reproduction among individuals explained by the environment (non-selective environmental variation), the genes (G), and their interactions (GxE), was modest compared to the magnitude of the non-selective demographic (i.e. stochastic) variation. We use these results to illustrate the influence that chance among individuals has on ecological and evolutionary population dynamics. We briefly discuss the advantages of our approach over alternative approaches, such as mixed effect models, that do not directly estimate or link to such demographic parameters. Our results highlight the challenge of distinguishing between adaptive genetic variability and neutral variation in evolutionary ecological, population biology, and demographic studies in the field.

## Materials and methods

*Plantago lanceolata* is a widely distributed short-lived perennial herb. In the experimental field site, located at the Shadwell Preserve of the Jefferson Monticello Foundation, Shadwell Virginia, USA, this species maintains a basal rosette of 1 to ~240 leaves (mean 12.5 ±11.53 SD). It germinates both fall and spring and remains green all year, thus individuals can be easily followed for size (number of leaves) and survival throughout the year. Seeds for this study were produced from crosses with parental individuals that had been randomly collected from the 70m * 35m field site. Parental genotypes were crossed using a modified North Carolina II design (Comstock and Robinson 1948). Here we used crosses that consisted of four sires crossed to each of two dams resulting in eight sire-dam combinations and 200 offspring from each. This was repeated for five unique sets of sires and dams (20 sires, 10 dams), 40 sire-dam combinations and 8000 individual offspring. In the analysis reported here, we used individuals from the 32 crosses with the largest replication (Roach et al. 2009). This design was used for cohorts 1 and 2 (planted in years 2000 and 2001) and one-half of the total number of individuals per cross was used for cohorts 3 and 4 (both planted in year 2002, spring and fall respectively, and for this analysis they were classified in the same age class). All cohorts had the same genetic structure when initially planted. Further details of the planting design and protocol are reported elsewhere (Roach et al. 2009). The analysis reported here began with the data collected in 2003 when 6913 individuals from these 32 crosses had survived. At this time, the number of individuals per cross, across cohorts, ranged from 283 to 376, and spatially these individuals were distributed across 17 randomized blocks and 64 plots. The matrix models constructed here use size (number of leaves) and survival data (censused and recorded in May/June each year) from 2003-2008. In total, there were 14082 events where size and survival in consecutive years was known. Data collected prior to 2003 was excluded because not all cohorts had been established before this period.

### Size structured matrix models

The analyses are based on discrete time (Markovian) size-structured matrix population models. Vital rates (growth, survival and fertility), were estimated using regression models (see below) as is normally done for integral projection models (IPM) (Ellner and Rees 2006), except that here we used the regressions to parameterize discrete size structured matrix models (Coulson et al. 2010). The stage structure consisted of 100 size classes (one for each number of leaves between 1 and 100+).

For each year-sire combination (6 years [2003-2008] and 16 sires, each sire crossed with two dams) we fit one model with size as the stage characteristic (Online Appendix Fig. S1). From each of these year-sire specific models we directly computed: population growth rate λ, generation time T_c_, net reproductive rate R_0_ (number of seedlings produced), expected reproduction (expected number of inflorescences produced) and variance in reproduction (variance in number of inflorescences produced) among year-sire individuals within the year-sire combination (non-selective demographic variation in reproduction), and life expectancy and variance in lifespan among year-sire individuals (non-selective demographic variation in survival within the year-sire combination) (Steiner et al. 2012, 2014) (Table 1).

**Table 1:** Notation and Equations. Details and proofs of equations are found elsewhere (Steiner et al. 2012, 2014)

| Description | Equation | Notes |
| --- | --- | --- |
| | $e_t$ | Vector of zeros with a 1 at position $t$ |
| | $e^T$ | Vector of ones, superscript $T$ denote transpose |
| Identity matrix | $\mathbf{I}$ | |
| Stage transition matrix | $\mathbf{P}$ | Includes survival and stage changes |
| Stage duration matrix | $\mathbf{N} = (\mathbf{I} - \mathbf{P})^{-1}$ | Elements quantify the expected time spent in each stage conditional on the birth stage |
| Mean Lifespan | $exL = e^T * \mathbf{I} * \mathbf{N} * e_t$ | |
| | $exLsq = e^T * \mathbf{I} * (2\mathbf{N} - \mathbf{I}) * \mathbf{I} * \mathbf{N} * e_t$ | |
| Variance in lifespan | $VarL = exLsq - (exL)^2$ | |
| Fertility matrix | $\mathbf{F}$ | |
| | $\hat{\mathbf{F}} = diag(\mathbf{F})$ | Diagonal elements of fertility matrix |
| Expected reproduction | $exR = e_t^T * \mathbf{F} * \mathbf{N} * e_t$ | |
| | $exRsqr = e_t^T * \mathbf{F} * (2\mathbf{N} - \mathbf{I}) * \hat{\mathbf{F}} * \mathbf{N} * e_t$ | |
| Variance in reproduction | $VarexR = exRsqr - (exR)^2$ | |
| Population growth rate | $\lambda$ =dominant Eigenvalue of $\mathbf{X}$, with population projection matrix $\mathbf{X} = (\mathbf{F} + \mathbf{P})$ | |
| Cohort generation matrix | $\mathbf{A}_c = \mathbf{F} * \mathbf{N}$ | |
| Net reproductive rate | $R_0 = \text{dominant Eigenvalue of } \mathbf{A}_c$ | |
| Right eigenvector corresponding to dominant eigenvalue of $\mathbf{A}_c$ | $cv$, normalized so to sum of components=1 | |
| Left eigenvector corresponding to dominant eigenvalue of $\mathbf{A}_c$ | $dv$, normalized so to $(dv, cv) = 1$ | |
| Cohort generation time | $T_c = (dv^T * \mathbf{N} * cv)/(dv^T * cv)$ | |

### Data analysis

For each model (i.e., year-sire combination), four regression functions were fit describing the relationship between the individual’s a) current size and survival, i.e. size-specific survival function; b) current size and size at time t+1, i.e. size-specific expected growth and the variance in size-specific growth among individuals; c) current size and reproduction; i.e. the size-specific probability of producing at least one inflorescence; and d) current size and number of inflorescences given that at least one inflorescence was produced, i.e. size-specific production of inflorescences (Online Appendix Fig. S1). In the final two years (2007 and 2008) one sire had too few surviving individuals (N=24 and N=9) to fit meaningful functions; we thus excluded these two years for this sire (number 16), which left us with 94 models (16 sires * 6 focal years; equals 96 minus the two years for sire #16). To estimate the non-selective environmental variance across all years and the additive genetic variance (G) across all sires, we used the demographic parameters from these 94 year-sire specific models weighted according to the number of individuals in each year-sire combination, i.e. year-sire combinations that had more individuals contributed more to the weighted variances across all years or sires than those with fewer individuals.

All functions were fit to square root transformed size measures to improve normality in residuals. For the survival and the reproduction functions, a binomial model with a linear and quadratic term for size was used (see example Fig. S1a and c). A Poisson distributed model structure with a linear and quadratic term for size was used for the number of inflorescences (see example Fig. S1d), and for the growth function a Gaussian linear model was used (see example Fig. S1b). It should be noted that around these size-specific functions there is substantial variance in the realized individual outcomes and among the year-sire specific functions. It is these two sources of variance that are core to our approach to decompose variance contributions. Because of the limited number of very large individuals, we binned all individuals ≥100 leaves (per year-sire combination) into one size class, and binned all individuals with ≥30 inflorescences (per year-sire combination) into one class. When estimating function parameters we included the dams as a random (intercept) effect to account for potential differences among dams.

To build full population matrixes ***X****gy*, for sire (*g*) in year (*y*), we made assumptions regarding recruitment. The field experiment did not allow direct recruitment of seedlings into the study site, thus did not have sire (*g*) specific recruitment rates *x*_*yg*_(*k,z*) where an individual of size (*z*) in year (*y*) contributes to the recruited individuals at size (stage) *k* at time t+1. For recruitment we used probability estimates from a separate study of seeds planted directly into the same field (Shefferson & Roach 2012). The probability of seedling establishment (0.1035) was computed from the germination rate of 0.69 and the seed to seedling survival rate of 0.15 (see robustness to assumptions about seedling establishment Online Appendix Table S1, Fig. S3). To account for differences among years in number of seeds produced per inflorescence, we took the mean number of seeds counted from approximately 150 randomly selected inflorescences each year. Mean number of seeds per inflorescence (±1 SE): year 2003, 70.72±2.83; year 2004, 52.83±2.69; year 2005, 42.5±2.43; year 2006, 42.07±2.28; year 2007, 50.86±1.82; year 2008, 27.84±2.09. For the seed to seedling size distribution, we assumed that these were equal across years and sires.

To test for potential effects of small-scale environmental differences throughout the 70m * 35m experimental field we corrected for variance among 17 blocks, small-scale geographic units across the experimental field. In this set of analyses we included one additional random effect, the block (N=17), for estimating the size dependent regression functions for each model (year-sire combination). Comparing the two sets of analyses (with and without correcting for variance associated to the blocks) provided insights on micro-climatic and small-scale environmental effects throughout the experimental field.

The four regression function parameters (Fig. S1) together with the number of seeds per inflorescence, the probability of seedling establishment, and the seedling size distribution were used to compute two 100*100 matrixes (100 size classes) for each year-sire combination. One matrix, the transition matrix (Table 1) contained survival rates and expected growth and the variance around that expected growth, i.e. the probability of transitioning from a given size at time *t* to size at time *t*+*1*. The column sums of this matrix *P* are smaller than 1 and are equal to the survival rates of individuals at a given size at time *t*. The other matrix, the fertility matrix *F* (Table 1), contained size-specific reproduction and recruitment rates, i.e. the probability of reproducing inflorescences with a yearly specific number of seeds, that then yields the recruitment of seedlings into the population depending on the current size at time *t*. We limited the maximum survival probability of any size to 0.95 (the results are robust to this assumption). The computation of the functions and the matrixes were done in program R (R Development Core Team 2012).

Each pair of matrices was used to compute: i) population growth rate λ, ii) cohort generation time, *T*_*c*_, iii) net reproductive rate *R*_0_(expected number of seedlings recruited), iv) mean lifespan, v) variance in lifespan, vi) mean reproduction (number of inflorescences), and vii) variance in reproduction for each year-sire combination (Steiner et al. 2012, 2014) (Table 1). The stochastic properties of these matrix models, which are rooted in Markov chain theories, allowed us to directly compute the expected variances in survival and reproduction. The duration matrix, *N*, (a.k.a. the fundamental matrix, see Caswell (2009)), was used to estimate the mean and variance in lifespan and reproduction for each year-sire combination (Steiner and Tuljapurkar 2012). The elements of this matrix quantify the expected time an individual spends in each stage conditional on the individual’s birth stage (Table1) (Steiner and Tuljapurkar 2012; Steiner et al. 2014).

Our models are based on year and sire, not age or cohort, because we found that size distributions did not shift along with age structure during the study (Online Appendix Fig. S2). Models that included only age and year, or age and sires (not shown) did not lead to qualitatively different results. Previous studies using this data (Roach and Gampe 2004; Roach et al. 2009; Roach 2012; Shefferson and Roach 2012) revealed that survival dropped significantly in 2003 for all cohorts or ages (Roach et al. 2009). A model comprising age-and-year-and-sire combinations could not be fit here because of lack of sample size; moreover, such a complex model would be difficult to interpret biologically.

We use matrix models to decompose the variability in survival and reproduction among individuals, which is in contrast to mixed effect models, GLMs, ANOVA, or similar approaches that are often used for a decomposition of the total variances, into the genetics, the environment (year), G × E, and the unexplained residual components (van de Pol and Wright 2009). These latter approaches produce biased estimates for various reasons. First, when estimates of genetic and environmental variance are based on low numbers of individuals within the same year or individuals that are closely related, such models are anticonservative and underestimate the stochastic component (Hadfield et al. 2010). Our analyses are less affected by such effects because of larger within year-sire numbers. Second, for a multiple cohort study such as ours with fewer cohorts than age classes, a mixed effect model would underestimate variance in mortality because of the limited age structure in a given year, which is a general problem in many studies tracking known-aged marked individuals, whereas variance in reproduction would be more accurately estimated (see Online Appendix Table S2). Third, a multitude of mixed models can be fitted with results that are often highly sensitive to factor combinations (see Online Appendix Table S2).

Our matrix models include aspects of mixed effect models but we do not include individual as random effects because our objective was to combine information about different life history traits and demographic rates. Our approach provides a direct link between survival probabilities and growth trajectories and we can easily combine different distributions (e.g. binomial for the probability to reproduce, Poisson for the number of inflorescences). The flexibility to combine information across models (e.g. survival and reproduction) is particularly challenging when using mixed effect models. Our approach made it possible to estimate, for each year-sire combination, a size structured matrix model that provides direct calculation of the non-selective demographic variation for each model (Steiner and Tuljapurkar 2012; Steiner et al. 2012, 2014).

## Results

The overall population growth rate λ was high, at 7.61, the cohort generation time T_c_ was 2.78 years, the reproductive value R_0_ was 20.0 expected seedlings recruited, the average lifespan was 2.67 years, the variance in lifespan was 6.32 years^2^, the average lifetime reproduction was 0.421 inflorescences, and the variance in reproduction was 1.42 inflorescences^2^. Our findings are consistent with previous studies with this data (Roach and Gampe 2004; Roach et al. 2009; Roach 2012; Shefferson and Roach 2012).

### Variance decomposition of lifespan and reproduction

Contributions to the total variance in lifespan and reproduction among individuals showed that only a small fraction (~0.5−1%) is explained by additive genetic (sire [G]) effects (Table 2). Non-selective environmental variability among years explained little variance (2.5-4.6%) in reproduction and about 25% of the variance in lifespan. A small fraction of the variation is explained by the genotype-by-environment (year-sire [G*E]) interactions (4.6% to 6.7%) (see also (Shefferson and Roach 2012)). The largest fraction of the variance is associated with unexplained variability in size within years and with sires that is not related to either size-specific reproduction or survival. Hence we argue that most of the variation in survival and reproduction was caused by non-selective demographic processes.

**Table 2:** Variance and the variance decomposition into genetics (sire), environment (variability among years), gene x environment interactions, and non-selective stochastic demographic variation for lifespan, and reproduction for two sets of models, i) overall model without accounting for any spatial environmental variability, ii) accounting for 17 blocks of the study field.

|  | Overall model |  | Block corrected |  |
| --- | --- | --- | --- | --- |
|  | Lifespan<br>(years) | Reprod.<br>(inflorescence) | Lifespan(years) | Reprod.<br>(inflorescence) |
| Absolute variances | 6.32 | 1.42 | 6.87 | 2.83 |
| Fractions of the variance decomposition |  |  |  |  |
| Genetics (sire) | 0.008 | 0.011 | 0.0075 | 0.0045 |
| Environment (year) | 0.245 | 0.046 | 0.250 | 0.025 |
| GxE | 0.067 | 0.064 | 0.065 | 0.046 |
| Stochastic | 0.680 | 0.878 | 0.678 | 0.925 |

We initially focused on variability among years, but geographic variation, even at a small scale, might influence survival and/or reproduction. To test for small-scale spatial variation we included the 17 blocks as a random variable when estimating the regressions for each model. Counter to expectations, variability for lifespan slightly increased when accounting for spatial block differences (Table 2). Variability in reproduction doubled when accounting for block differences compared to the initial model with no spatial component (Table 2). Further, and again counter to expectations, there was no increase in genetic (sire) or environmental (among year) contribution after correcting for spatial environmental differences (Table 2). We also evaluated small-scale microsite variability using smaller units of 64 plots, but our results (not shown) were no different from the analysis with the 17 larger blocks.

Our study clearly shows the importance of variability among individuals within each genotype-by-environment (sire-by-year) class, which is a new finding from these data. These results show how key demographic parameters vary among models parameterized for different years and genotype combinations. Specifically, expected population growth rate λ varied substantially among years (Fig. 1A); generation time, T_c_, was relatively short in the first three years compared to the remaining years of the study (Fig. 1C), as was net reproductive rate R_0_ (Fig. 1E), average lifespan (Fig. 1G), and expected reproduction (Fig. 1I). Additionally, offspring of different sires differed substantially in population growth rate λ (Fig. 1B), generation time, T_c_, differed slightly less among sires (Fig. 1D), and net reproductive rate, R_0_ varied tremendously among sires (Fig. 1F). Lifespan varied to a lesser degree (Fig. 1H) compared to expected reproduction (Fig. 1J).

**Fig. 1.**
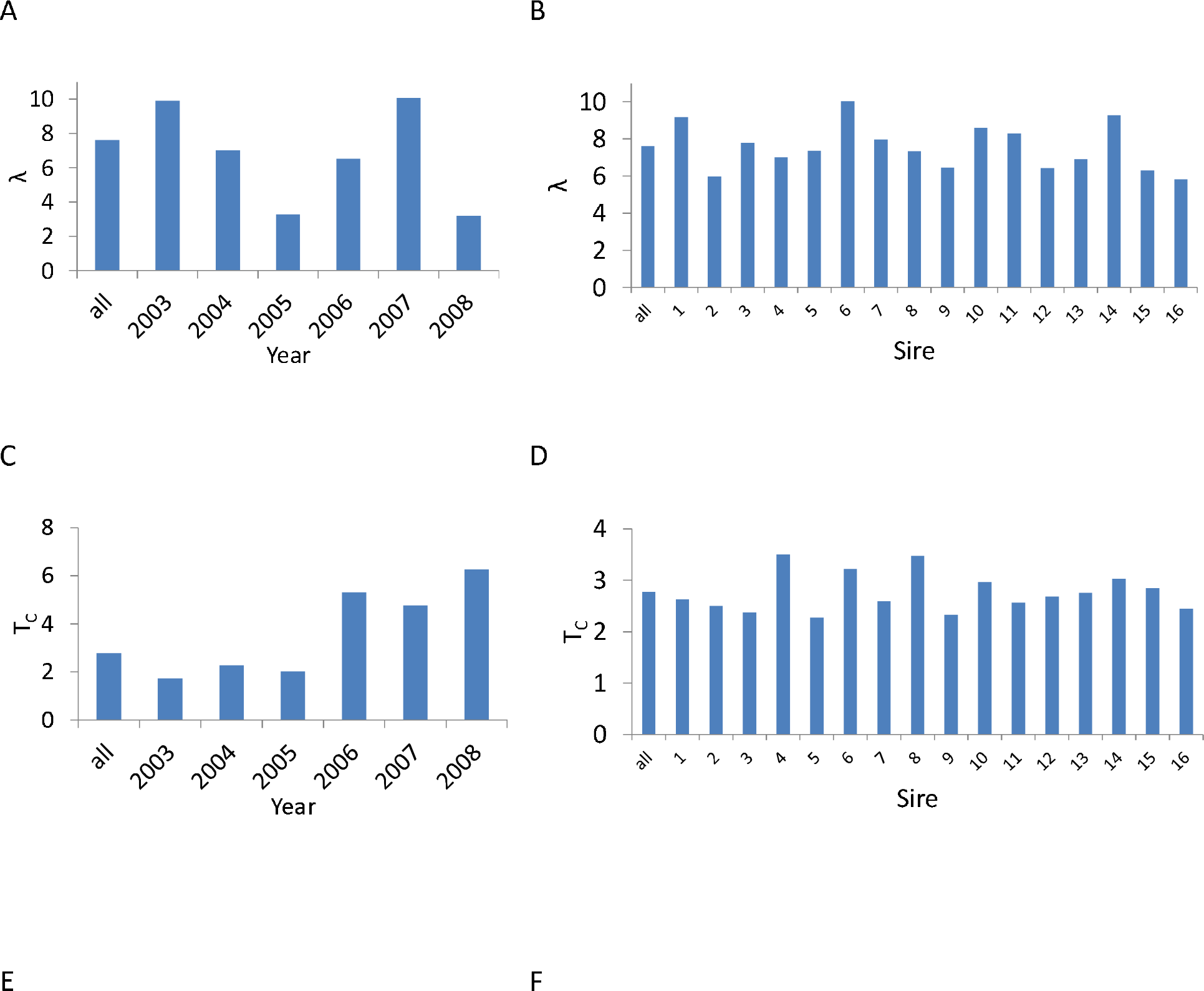
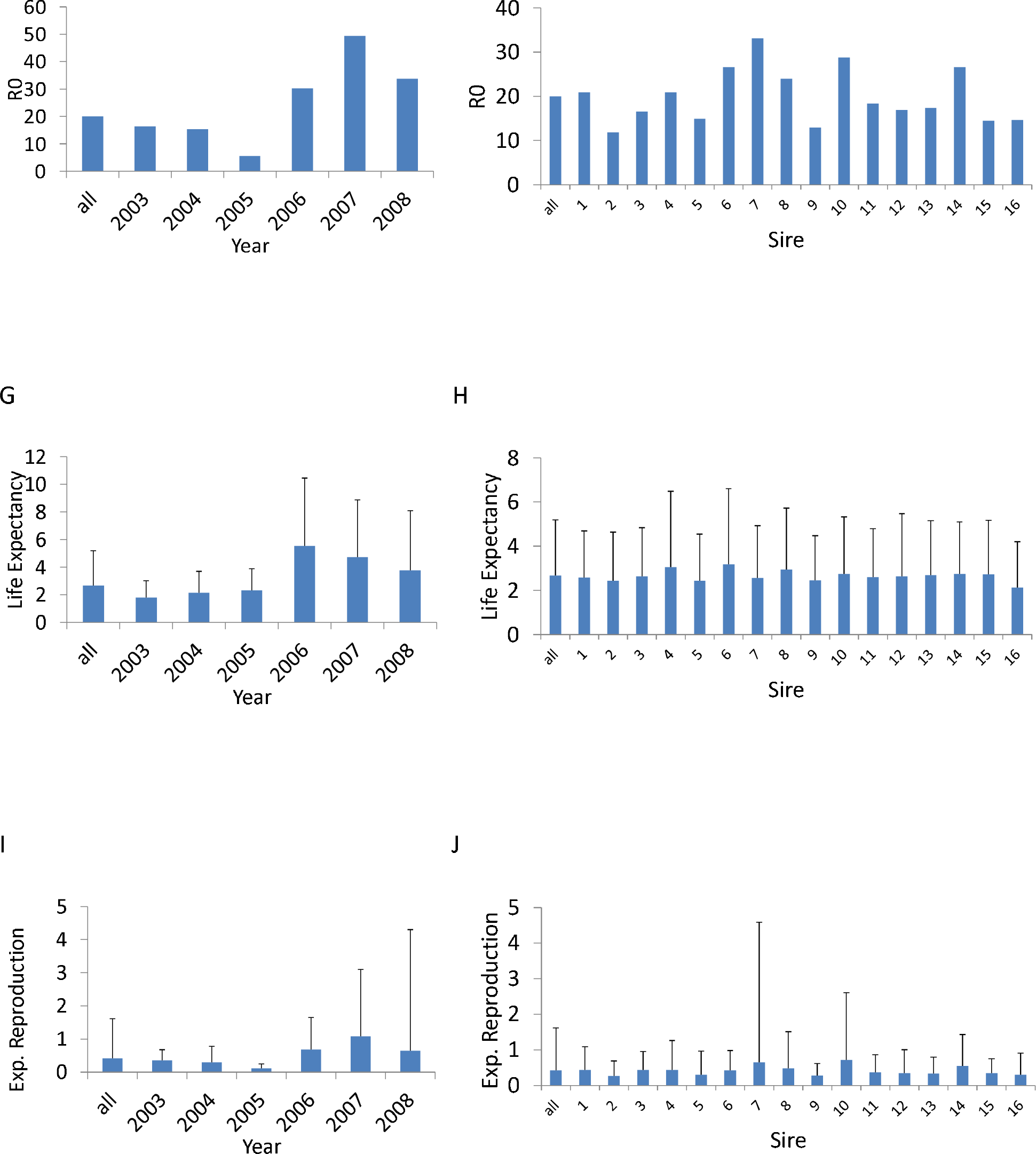
Differences in population growth rate λ (A, B), cohort generation time T_c_ (C, D), net reproductive rate R_0_ [expected number of seedlings recruited] (E, F), life expectancy (G, H), and expected reproduction [expected number of inflorescences] (I, J) among years (A, C, E, G, I) and sires (B, D, F, H, J). The most left bar depicts the value across all years or sires, weighted by the individuals within each year-sire combination. For life expectancy and reproduction (G, H, I, J) we plotted the mean + Stdev. This standard deviation comes from unexplained variability in size within years and sires that is not related to either reproduction or survival, we argue here that this variability is largely due to non-selective variation among individuals sharing the same sire and environment (year). See also Table 1, Appendix S1, (Steiner & Tuljapurkar 2012; Steiner *et al.* 2014).

## Discussion

We have used a plant dataset to illustrate the importance of non-selective demographic processes on population dynamics. We analyzed fitness components and overall fitness, λ, by using experimental data to parameterize stage-structured matrix population models for each distinct gene-environment combination. Our approach allowed us to examine genetic, environmental and stochastic variation in survival and reproduction. We found that selective variation, genetic (G) and genotype-environmental (GxE) variation makes a small contribution to total variance in survival and reproduction, as does non-selective environmental variation, whereas non-selective demographic variation is very large. The design and richness of this data set allowed us to extend previous analytical approaches (Coulson et al. 2008, 2010). The important features of the data that made our analyses possible included the detailed demographic information on many individuals stemming from a small number of genotypes (sires), and the negligible detection and measurement errors, for leaf counts, inflorescence counts and survival, for the marked and mapped individual plants. Additionally, the experimental design excluded within-species density dependence, because the spacing among the planted individuals was large enough to avoid direct competition. Natural recruitment from the experimental plants was not allowed thus there was no increase in within species density over time.

To critically examine our claim that the dominant source of variance in fitness components is non-selective demographic variability, we analyzed two alternative sources of variation, the micro-site environment and genetics. With respect to the environment, the randomized block planting design was used to minimize small-scale environmental influences in a field setting where spatial environmental differences among individuals cannot be completely avoided. If such differences contributed to major variation among individuals, we would expect non-selective stochastic variability in fitness components to decrease substantially after we accounted for small scale geographic differences (blocks) in our models, while the relative genetic (sire) and among-year environmental contributions would increase. But accounting for the blocks, or even for smaller spatial scale environmental differences of 64 sections of the 70 m * 35 m field site (results not shown), did not reduce the overall variability in fitness components as expected. Moreover, the relative genetic and among year environmental contributions did not systematically increase, as expected (Table 2). We cannot completely exclude that even finer scaled environmental differences might influence survival and reproduction among individuals within this population but without additional experimentation this source of variation cannot be clearly identified. Moreover, it is clear that non-selective demographic processes do contribute much of the variability in fitness components.

Genetic variability may also contribute to fitness variability among individuals in this non-clonal species. To include this source of variation, our primary analyses accounted for the effects of the dam. Another approach is to reverse the genetic focus by swapping dams and sires, i.e., assess dam effect while correcting for sires as random effects. When we did this analysis, our results do not change qualitatively (analysis not shown). Our low estimates of additive genetic (sire) and gene-by-environment contributions suggest that additional genetic differences among individuals stemming from the same sire cannot account for much of the unexplained variability in survival and reproduction.

Our models estimated a very high overall population growth rate λ of 7.42, which is obviously unrealistic for any natural population (see similar estimates from a deterministic life table response experiment (LTRE) analysis of the same experimental population in Shefferson and Roach (2012)). Because of the high accuracy in tracking individuals, their survival and the number of inflorescences, we believe this high estimate comes from ignoring losses in the transition from seed to established seedling. Seed predation (Lacey et al. 2003) and intraspecific density regulation were excluded in the experiments and would lower population growth. Irrespective of the exact cause for the high estimate of population growth rate, our main result that decomposed variability into genetic, environmental, and stochastic components should not be qualitatively affected (Online Appendix Fig. S3, Table S1).

Variability in the population growth rate λ among sires (Fig. 1B) is less extreme compared to the variability among environments (years; see Fig. 1A). The sires that would be expected to go extinct first (Fig. 1B, sire 2, 9, & 16), are the ones that have low reproductive rates (Fig. 1F) and tend to have fast life histories (Jones et al. 2008), i.e. low cohort generation time (Fig. 1D). However, these patterns require closer examination: Sire 7 has the highest expected reproduction, with a mean lifespan and a generation time that does not differ much from the population average. One would therefore expect sire 7 to have high fitness (λ) (Steiner et al. 2014), but this expectation does not hold because reproduction varies substantially within the sires’ offspring (Fig. 1J). These results highlight how influential the within year-sire variation in survival and reproduction are for evolutionary population dynamics and caution us not to interpret demographic parameters in isolation, at least when making evolutionary ecological predictions.

Despite the dominating influence of non-selective demographic effects, environmental variation among years was strong. During the course of the six years analyzed in this study, there were three years of high mortality, which suggests stressful environmental conditions (2003-2005) (Roach et al. 2009). The λ decreased during those three years (Fig. 1A). Generation time and net reproductive rate R_0_ indicate fast life histories with short expected lives and low expected lifetime reproduction (Fig. 1). Interestingly, there were no carryover effects of the stress in these years, with respect to survival and reproduction (see also little shift in size distribution across the years Online Appendix Fig. S2). This lack of carryover is particularly interesting given that small, and in particular small, old, individuals died at high rates during this stressful time (Roach 2012).

Our estimates of the relative size of additive genetic variation might be considered small, but a low additive genetic contribution to the total variability in survival and reproduction among individuals is not surprising from a population genetics perspective or basic evolutionary theory (Fisher 1930; Wright 1931; Crow and Kimura 1970). Indeed, many studies of natural populations find low estimates of heritabilities for fitness components (Merilä and Sheldon 1999, 2000, Kruuk et al. 2000, 2001; Sheldon et al. 2003; Teplitsky et al. 2009). Given low heritabilities, and the sample sizes (per genotypes) typically found in ecological studies, detecting any evolutionary change within the time scale of such studies will be challenging (Steiner and Tuljapurkar 2012). Our low estimates of additive genetic variation do not mean that the variability is evolutionary unimportant but rather, that selective changes will be slow and genetic drift enhanced, particularly in populations with long or complex life cycles (Endler 1986; Charlesworth 1994; Hartl and Clark 2007; Steiner and Tuljapurkar 2012).

Our estimates of the amount of non-selective demographic variation are consistent with what has been found for fitness components in individuals of model organisms raised in the lab under highly controlled conditions. Even among isogenic individuals under lab conditions the coefficient of variation (CV) of the stochastic demographic component ranges between 0.24 to 1.33 in lifespan (*Caenorhabditis elegans* 0.24-0.34 (Finch and Kirkwood 2000; Kirkwood et al. 2005), *Caenorhabditis briggsae* 0.31-0.51 (Schiemer 1982), *Saccharomyces cerevisiae* (0.37)(Kennedy 1994), *Escherichia coli* 0.4-0.6 (Steiner et al. 2017; Jouvet et al. 2018)). Less controlled studies that include individuals with more genetic variation do not differ much from these patterns in the CV for lifespan, for example in laboratory reared mice (0.19-0.71)(Finch and Kirkwood 2000) or *Drosophila melanogaster* 5.98-13.48 (Curtsinger et al. 1992). These values are in the range of the values we detect here: CV 0.96 for lifespan, and 3.97 for reproduction (non-block corrected estimates). Such estimates are also well within the range of variability in lifespan and reproduction of other natural populations, even though the decomposition into non-selective stochastic components is often not possible under less controlled conditions (Tuljapurkar et al. 2009; Steiner et al. 2010).

The large variability in fitness components among individuals that cannot be explained by genetics or the environment remains a mystery if not considered in the context of non-selective stochasticity. Other studies, outside the field of evolutionary ecology, that are at a more mechanistic level support the idea that stochastic events play a major role in determining outcomes at all levels, from stochastic gene expression, to the protein level, to the cell and organismal organization level (Elowitz et al. 2002; Kærn et al. 2005; Balázsi et al. 2011; Norman et al. 2015; Vera et al. 2016).

Our approach differs from previous models that estimate non-selective demographic variability (Engen et al. 2005; Vindenes et al. 2008) by including stages and not just age. In previous models, demographic variability has been estimated as the variance and covariance in survival and fertility within an age class, but these models do not provide a mechanism to correlate performance across ages because age classes are assumed to be independent. Further, these models do not consider cohorts, which then makes it impossible to compute life history traits such as an individuals’ age at death, or lifetime reproductive success. This limitation thus makes alternative models less applicable for assessing life history tradeoffs, such as the fundamental tradeoff between survival and reproduction (Stearns 1992). Additionally, previous models have included approximations for stochastic dynamics (transient dynamics), whereas the models presented here are deterministic. Excluding stage dynamics (here size as stage) would lead to very different estimates of non-selective demographic variability, because in our model the stage dynamic is one of the main processes generating variance and is crucial for the computation of the correlation between growth, survival and reproduction within and among ages (the latter through the Markovian structure of the model). Such stage dynamics thus allow us to compute the non-selective stochasticity we were mainly concerned with, and thereby account for the importance of quantitative trait dynamics for life histories. Our focus on individual measures also recognizes the central role that individuals play in demographic and population changes, including fundamental tradeoffs and covariances between longevity and lifetime reproduction (Stearns 1992; Metcalf and Pavard 2007). Here, we have shown that variation driven by stochastic demographic processes can be precisely quantified by estimating and analyzing a (stage or stage-age) structured matrix model, and this approach provides us with a better understanding of population dynamic processes, including the effect of stochastic events on population growth and other fitness related demographic parameters.

In ecology and conservation biology, the role of stochastic demographic processes has been mostly investigated for population extinction processes rather than evolutionary ecological processes (Fox and Kendall 2002; Kendall and Fox 2002; Lande et al. 2003)(but see Vindenes et al. 2008, Bolnick et al. 2011). Ecological models provide insight into the effects of stochastic environmental variation on vital rates, but surmise that stochastic demographic variation is of little importance because, as long as the populations are not very small, extinction is not influenced by such processes (Lande et al. 2003). However, in the context of evolutionary dynamics, the adaptive potential to respond to climate change can be substantially influenced by large amounts of stochastic demographic variation and may in the long run influence extinction even in large populations (Hartl and Clark 2007; Steiner and Tuljapurkar 2012). The data used here is a single illustrative example that shows that large within year-sire variation in reproduction does not lead to high expected growth rates of these sires. It highlights, that even in large populations, evolutionary population dynamics, and consequently long run extinction, can be influenced by such neutral variation.

## Conclusions

Historically, unexplained variation (residual error) has often been interpreted as a lack of knowledge of underlying causes, measurement error, and/or limited control of the conditions under which experiments are conducted. Recent studies have been devoted to determining the impact of stochastic events on variation at the molecular, cellular and organismal level. At each of these levels substantial evidence suggests that stochasticity plays an important role (Elowitz et al. 2002; Kærn et al. 2005; Balázsi et al. 2011; Norman et al. 2015; Vera et al. 2016). In some circumstances the cause of the “stochastic” outcome can be tracked down to a mechanistic cause at a lower level, but this mechanistic cause may be triggered by other stochastic events. If such cascading stochastic (snowball) events play a major role in determining demographic variation, then variability in fitness components has to be seen as neutral and not driven by the environmental or genetic variability among individuals. Controversial discussions about neutral theories, in molecular evolution or community ecology, also highlight the idea that these processes are ubiquitous, at many different levels, but are not easy to quantify (Ohta and Gillespie 1996; Alonso et al. 2006; Hughes 2008).

We have shown that large amounts of variability in fitness components among individuals in this study are likely due to stochastic demographic processes and such neutral variability has significant effects on population dynamics and demographic parameters. Our understanding of this type of variability and its impact on the evolution of phenotypic variability is limited and we call for more attention to and focus on understanding such variation. These neutral processes have ecological and evolutionary consequences, but neither our current theories nor our empirical understanding are sufficient to explain their evolution and maintenance.

## Acknowledgements

We were supported by Marie-Curie PIEF-GA-2009-235205, the Max-Planck Society, NIA P01-AG0225000-01 (UKS, ST) and NIH P01-AG8761 (DAR). We thank Tim Coulson, Johan Dahlgren, Dan Levitis, and Anne Charmantier for discussions and comments.

## Online Appendix: Examples of regression functions fit to parameterize each (year-sire specific) mixed effect model

**Fig. S1:**
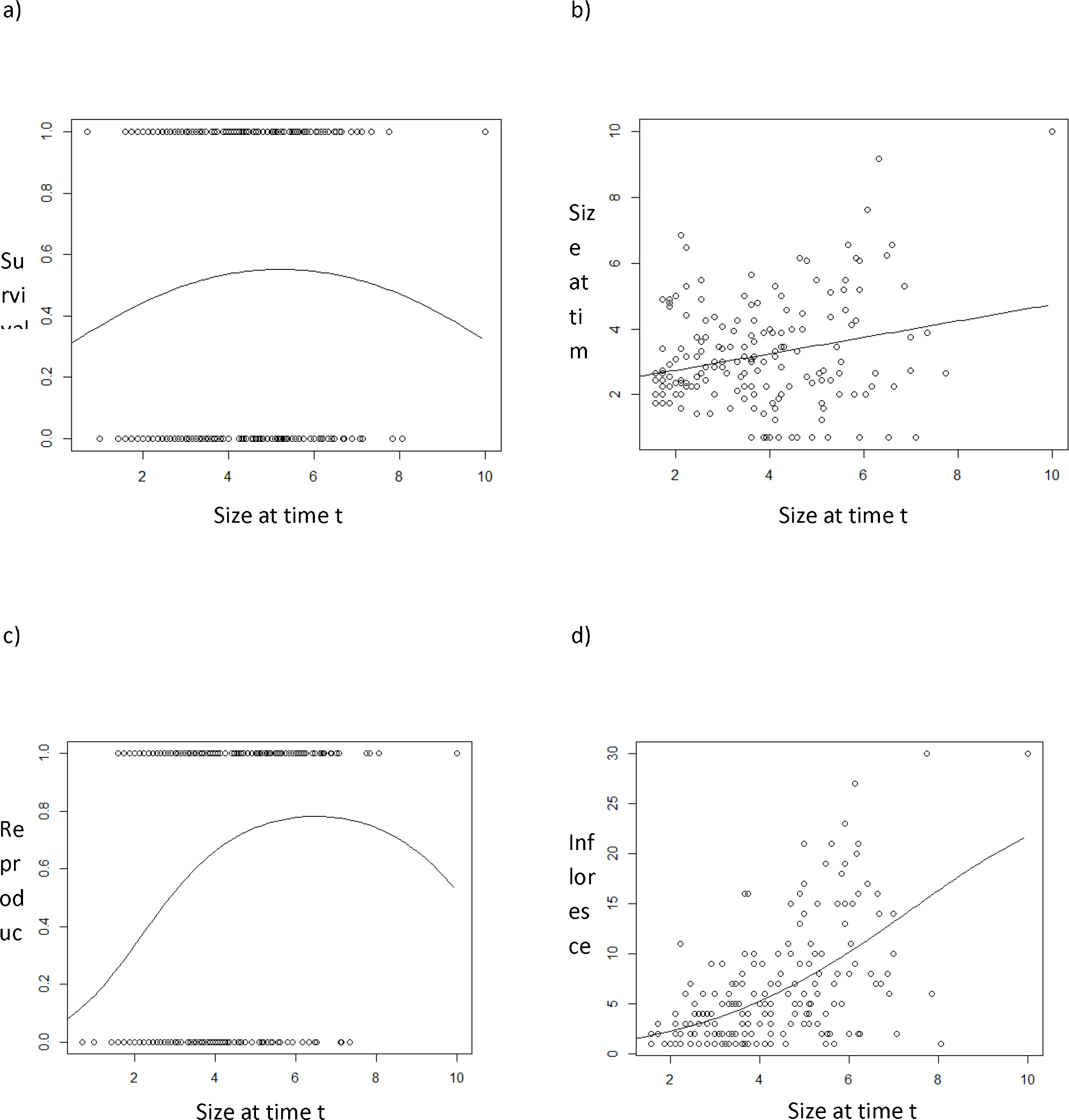
Examples of four regression functions that were fit to the initial data from the marked individuals for each model. These regressions describe the relationship between the current individual’s size and a) survival, b) size at time t+1, c) reproduction, i.e. the probability of producing at least one inflorescence given the current size, and d) number of inflorescences given that at least one inflorescence was produced. For the growth function b), the variance was estimated and used for computing the matrixes and the sources of non-selective demographic variation in lifespan and reproduction that was not related to size. The data and functions shown here as examples, are for sire 1, in year 2003 (sire-year combination). These four functions were used to parameterize the size structured matrix model, i.e. the two 100*100 matrixes. One model was fit for each year-sire combination. All size measures were square root transformed.

## Variation in the size distribution over the different study years.

The multiple cohort design of the experiment led to a shift in the age structure to older age classes during the study. Given that age and size might be partly correlated, the shift in the age structure might lead to a shift in the size structure with increasing years. Note that we excluded data collected prior to 2003 because not all cohorts had been established before this period. Fig. S2 illustrates that there was no systematic shift in size structure with shifting age structure in this study. There is variability in the size structure among years (mainly driven by the environmental variation), but a systematic shift in size distribution (expected towards larger sizes) during the experiment was not found. The first year, 2003 shows a distribution with relatively large individuals, 2004, 2005, and 2006, show distributions with relatively small individuals, while 2007 and 2008 again show slightly larger individuals.

**Fig. S2:**
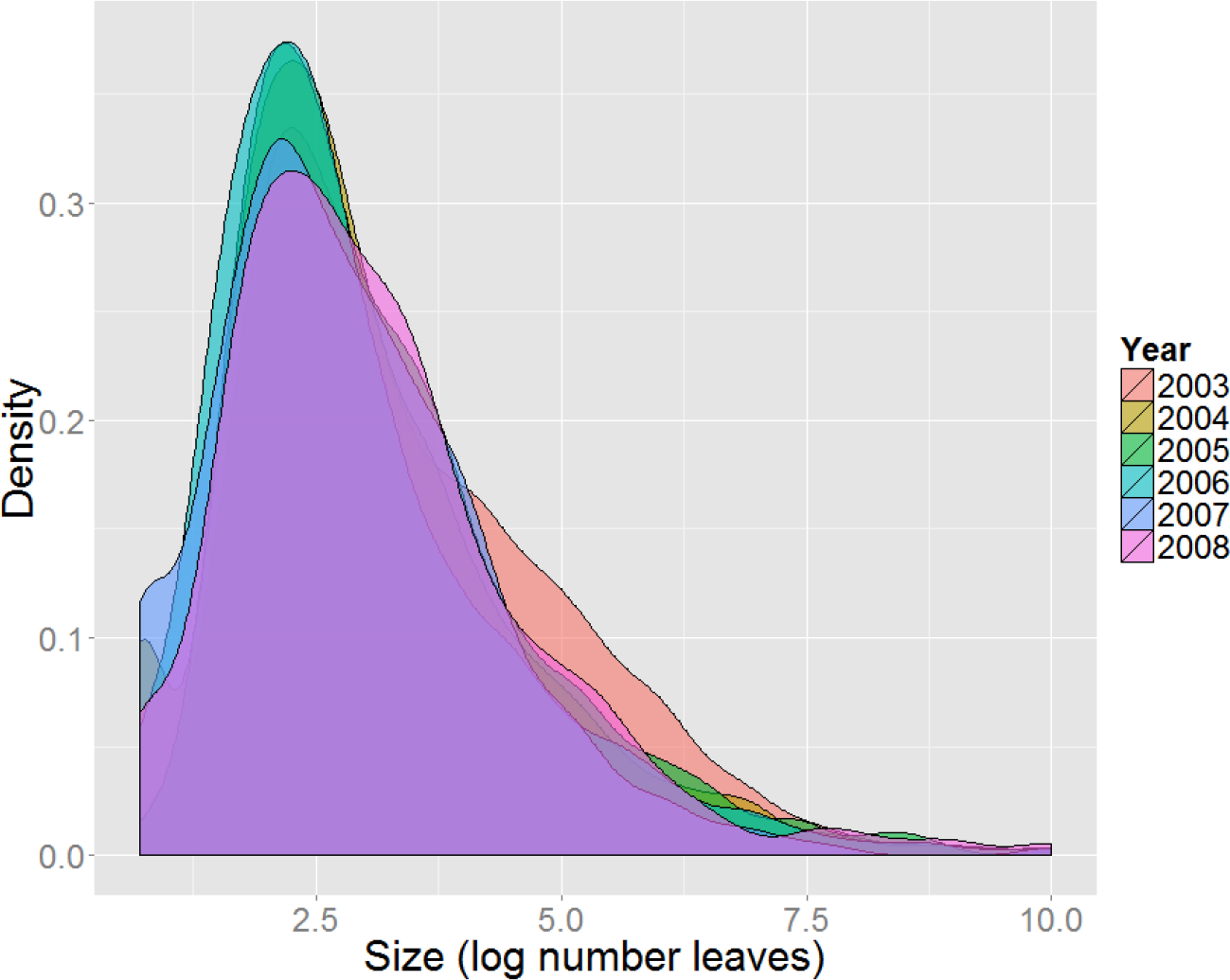
Size distribution (log transformed, number of leaves) for the six study years, 2003-2008.

## Robustness of results to assumptions about seedling establishment

**Table S1:** Testing for robustness of the main result (stage-structured matrix models; Table 2 main text) to changes in the probability of seedling establishment. Here we only present results without accounting for any spatial environmental variability (i.e. no block effect included). In the original analysis (Table 2) seedling establishment was estimated at 0.1035, here we lowered the seedling establishment to 0.01035, hence assume a ten times lower germination rate (0.069) or seed to seedling survival rate (0.015) compared to the estimates used in the main test where seedling establishment was based on a separate study that planted seeds directly into the field (Shefferson & Roach 2012). All other parameters and estimation have been kept exactly the same.

|  | Model with lowered seedling establishment |  | Original model (Table 2)<br>Seedling establishment (0.1035) |  |
| --- | --- | --- | --- | --- |
|  | Lifespan<br>(years) | Reprod.<br>(inflorescence) | Lifespan<br>(years) | Reprod.<br>(inflorescence) |
| Absolute variances | 6.32 | 0.0285 | 6.32 | 1.42 |
| Fractions of the variance decomposition |  |  |  |  |
| Genetics (sire) | 0.008 | 0.007 | 0.008 | 0.011 |
| Environment (year) | 0.245 | 0.024 | 0.245 | 0.046 |
| GxE | 0.067 | 0.047 | 0.067 | 0.064 |
| Stochastic | 0.680 | 0.922 | 0.680 | 0.878 |

Reproduction does not influence survivorship and lifespan, for that the variance decomposition for that part of the analysis does not change. However, reduced seedling establishment lowered the absolute variance explained by reproduction, reduced the genetic, environmental and gene*environmental contribution to the total variance, and increased the stochastic component. The overall quantitative pattern remained similar. Further details of the effect on reduced seedling establishment are illustrated in Fig. S3.

**Fig. S3:**
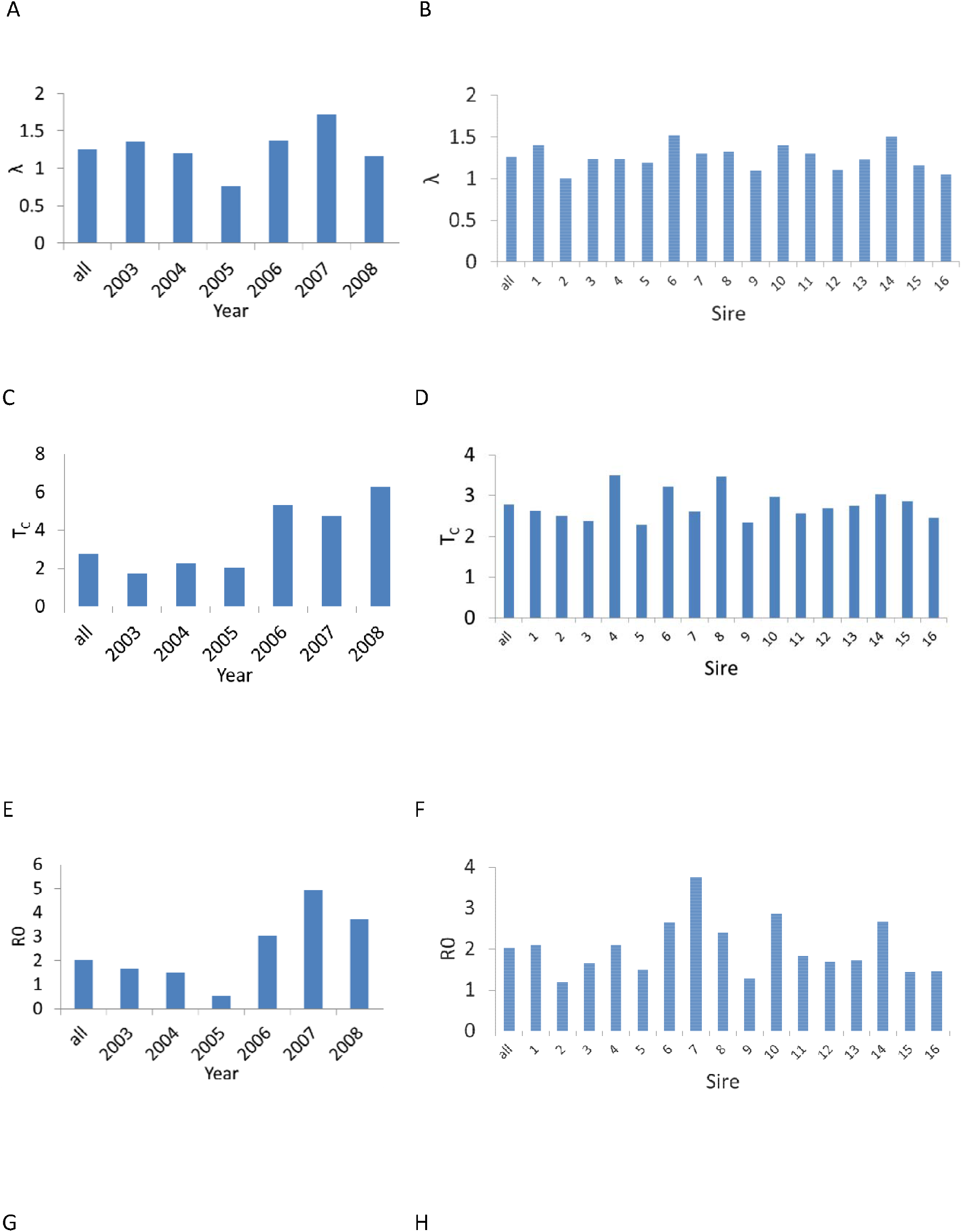
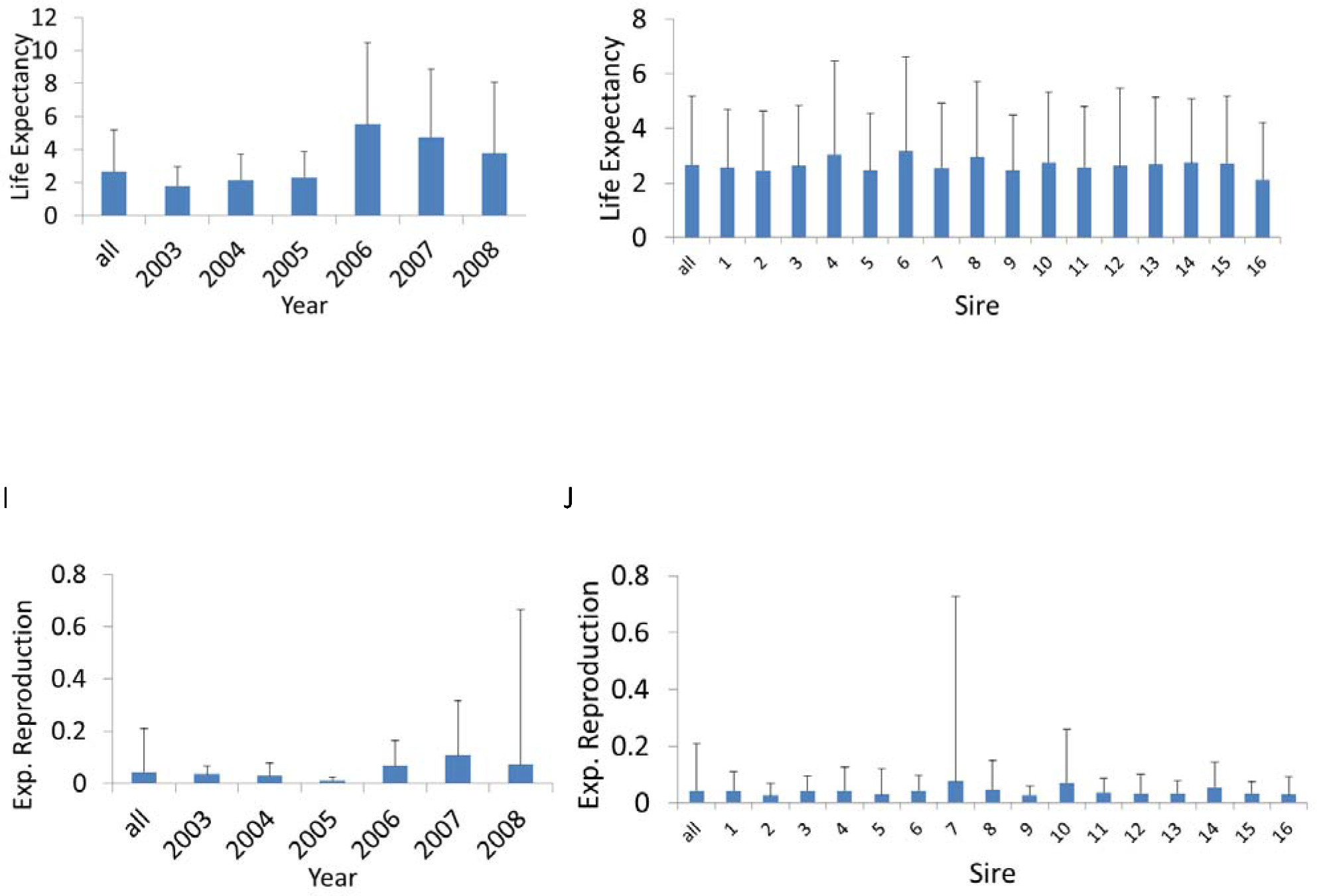
Differences in population growth rate λ (A, B), cohort generation time T_c_ (C, D), reproductive value R_0_ [number seedlings recruited] (E, F), life expectancy (G, H), and expected reproduction (I, J) among years (A, C, E, G, I) and sires (B, D, F, H, J). The most left bar depicts the value across all years or sires. For life expectancy and reproduction (G, H, I, J) we plotted the mean + Stdev. The seedling establishment is lower (0.01035) compared to the original analysis presented in the main text (Fig. 1), all else is kept constant.

## Generalized linear models and mixed effect models

Alternative approaches to our matrix model approach include GLMs and mixed effect models (GLMMs). We present a set of such models, even though they cannot be directly compared because they include different information. A multitude of linear models could be fit and our limited selection presented here is based on fitting models that include factors similar to the matrix model we have used in the main text. For the GLM and GLMMs we fit survival (binomial variable) and the number of inflorescences (Poisson error distribution) as the response variable respectively. For the mixed effect model we included sire and year, their interactions, as well as their interactions with (log)size as fixed effects, and included dam as random (intercept) effect. For a variance decomposition, we estimated the total variance explained by the fixed (and random effects) using the method developed by Nakagawa, & Schielzeth (Nakagawa and Schielzeth 2013), function r.squaredGLMM, program R package {MuMIn}, that estimates a R^2^ value for all fixed effects together and the combined R^2^ of fixed and random effects. The residual variance (1− R^2^) is then interpreted as unexplained variance (non-selective demographic variance and non-selective noise). The ANOVA command in program R was then used to decompose the total variance explained by the fixed and random effects (R^2^) into the specific fixed effects (G, E, GxE). Similar sets of models were fit for the GLM’s except that the dam was included as an additional interacting effect instead of a random effect. Because the random dam effect was estimated to explain only a very small fraction of the variance, a third set of models were setup where we dropped the dam effect completely. Despite various convergence problems, we present the results in Table S2, but raise caution for interpretation. All models were fit in program R (R Development Core Team 2012).

**Table S2:** Variance decomposition into genetics (sire), environment (variability among years), gene*environment interactions, and stochastic variation for survival, and reproduction (% inflorescences) of three sets of models, i) GLMM with dam as random (intercept) effect, ii) GLM with dam as interactive fixed effect, iii) GLM without any dam effect.

|  | Mixed effect model (GLMM) |  | GLM (including interactions with dam as fixed effect) |  | GLM (without any dam effect) |  |
| --- | --- | --- | --- | --- | --- | --- |
|  | Survival | Reprod. (# inflorescence) | Survival | Reprod. (# inflorescence) | Survival | Reprod. (# inflorescence) |
| Fractions of the variance decomposition |  |  |  |  |  |  |
| Genetics (sire) | 0.109 | 0.3228 | 0.0056 | 0.012 | 0.037 | 0.280 |
| Environment (year) | 0.160 | 0.0437 | 0.0996 | 0.351 | 0.062 | 0.073 |
| GxE | 0.052 | 0.0144 | 0.0198 | 0.022 | 0.012 | 0.015 |
| Random effect (dam effect) | 0.001 | * | - | - | - | - |
| Residual error | 0.677 | 0.619* | 0.8750 | 0.615 | 0.889 | 0.632 |

\*Convergence problem for r.squaredGLMM, hence residual variance estimated by getME(object, "devcomp"), R package lme4.

Table S2 illustrates the difficulty in determining what factors should be included and how sensitive such models are to different parameter combinations. The variance in survival explained by the fixed effects drops dramatically when the dams are not included as a random effect, even though the random effect in the GLMM accounts only for a very negligible amount of variation (0.001). The variance decomposition in survival changes between GLM with and without dam as fixed (interactive) effects and the additive genetic contribution changes markedly from 0.56% to 3.7%. Residual variance for reproduction does not change as dramatically between GLMM and GLM’s, though the contributions of the different fixed effects changes extremely between GLMM and GLM’s. In the GLM with dam as fixed (interactive) effects, 35% of the total variance is explained by the environment (year) and only little is explained by the genetics (~1%). When we dropped the dam effect from the model or fit a GLMM with dam as random effect, most of the variance is explained by the genetics (28% and 32% respectively), and only relatively little (7.3% or 4,4%) is explained by the environment (year).

It should be noted that the variance decomposition (Table S2) could not be directly compared to the matrix model results in the main text (Table 2), because the matrix models decomposed variance in longevity, including the size and growth functions, whereas in the linear models (GLMM & GLM) survival (binomial variable) was decomposed. For reproduction in the matrix models, we decomposed variance in reproduction that incorporated both probability of reproducing (binomial variable) as well as the number of inflorescences and their relationship to size, while in the linear models (GLMM & GLM, Table S2) only number of inflorescences was included.

